# Phage display enables machine learning discovery of cancer antigen specific TCRs

**DOI:** 10.1101/2024.06.27.600973

**Authors:** Giancarlo Croce, Rachid Lani, Delphine Tardivon, Sara Bobisse, Mariastella de Tiani, Maiia Bragina, Marta AS Perez, Julien Schmidt, Philippe Guillame, Vincent Zoete, Alexandre Harari, Nathalie Rufer, Michael Hebeisen, Steven M Dunn, David Gfeller

## Abstract

T cells targeting epitopes in infectious diseases or cancer play a central role in spontaneous and therapy-induced immune responses. T-cell epitope recognition is mediated by the binding of the T-Cell Receptor (TCR) and TCRs recognizing clinically relevant epitopes are promising for T-cell based therapies. Starting from one of the few known TCRs targeting the cancer-testis antigen NY-ESO-1_157–165_ epitope, we built large phage display libraries of TCRs with randomized Complementary Determining Region 3 of the β chain. The TCR libraries were panned against the NY-ESO-1 epitope, which enabled us to collect thousands of epitope-specific TCR sequences. We then trained a machine learning TCR-epitope interaction predictor with this data and could identify several epitope-specific TCRs directly from TCR repertoires. Cellular binding and functional assays revealed that the predicted TCRs displayed activity towards the NY-ESO-1 epitope and no detectable cross-reactivity with self-peptides. Overall, our work demonstrates how display technologies combined with machine learning models of TCR-epitope recognition can effectively leverage large TCR repertoires for TCR discovery.

## Introduction

T cells play a key role in infectious diseases and cancer immunotherapy ^1–3^. The T-cell response is initiated by the binding of T-Cell Receptors (TCRs) to specific peptides (referred to as epitopes) displayed on the surface of cells by Major Histocompatibility Complex (MHC) molecules (also called Human Leukocyte Antigens or HLA). TCRs are heterodimer surface proteins composed of an α and a β chain. TCRs show extensive sequence diversity across different T cells and approximately 10^11^ T cells with distinct T-Cell Receptors are constantly circulating in the human body ^4–6^. The TCR sequence diversity is achieved during the V(D)J recombination where a unique combination of the germline-encoded V and J segments, respectively V, D and J segments, are selected to form the α, respectively β, chain. Both chains undergo additional nucleotide insertions and deletions at the V(D)J junctions, thereby further increasing the diversity of TCR sequences. The V segments contain two complementarity-determining regions (CDR1 and CDR2) which primarily mediate contact with the MHC, and a third one (CDR3, located at V(D)J junctions) which is mainly involved in recognition of the epitope.

TCRs recognizing clinically relevant epitopes represent promising therapeutic agents for T-cell based immunotherapy. For instance, T cells enriched in TCRs recognizing cancer epitopes have been infused into patients to mount responses against different malignancies ^7,8^. TCRs recognizing specific epitopes also show promise for diagnostics since their presence in the TCR repertoire of a patient can inform clinicians of the past or present immunological status of this patient ^9–12^. From a more fundamental point of view, epitope-specific TCRs provide key information to characterize the specificity of TCR-epitope interactions ^13,14^.

Binding and activation assays have been widely used to isolate and sequence epitope-specific TCRs ^15,16^. These approaches typically involve *in vitro* stimulation of primary T cells from donors with the epitope of interest, followed by isolation and TCR-sequencing of the epitope-specific T cells. Binding assays use individual peptide-MHC (pMHC) multimers ^17–19^ or multiplexed DNA barcoded pMHC multimers ^20,21^, coupled with flow cytometry to isolate epitope-specific T cells. Functional assays use specific markers, such as CD137, PD-1 or CD69, to identify epitope-specific T cells which are activated by epitope stimulation ^22,23^. These approaches have enabled researchers to sequence thousands of epitope-specific TCRs for several immunodominant epitopes restricted to frequent MHC alleles ^10,24–26^. However, the number of epitopes with enough TCRs for in-depth characterization of their specificity is still limited. For instance, only 27 epitopes have more than 100 known αβTCRs in public databases^10^. The scarcity of data is especially pronounced for cancer epitopes, which are more challenging to profile in standard binding or functional T-cell assays since the TCR repertoire of patients or donors typically contains only very few (if any) TCRs recognizing such epitopes ^27^. A prototypical example is the HLA-A*02:01 restricted NY-ESO-1_157–165_ epitope ^28–30^ (hereafter referred to as NY-ESO-1). NY-ESO-1 is a widely studied cancer-testis antigen with very low expression in normal, non-germline tissue, but it is aberrantly expressed in many tumors ^31,32^. NY-ESO-1 can elicit a T cell response and therefore represents a promising target for many T-cell based immunotherapies ^28–30^. NY-ESO-1 reactive T cells have been reported in the blood of some patients with metastatic melanoma ^33,34^, but are rare in other patients or in healthy donors. Currently less than fifteen naturally occurring NY-ESO-1 specific TCRs are available in databases of epitope-specific TCRs like VDJdb ^27^. Such low numbers make it challenging to characterize the specificity of TCRs recognizing this epitope ^35–37^.

Naturally occurring TCRs targeting NY-ESO-1 are usually of low affinity (around 10 μM) ^38–40^ and several approaches have been used to design affinity-enhanced TCRs for therapeutic applications. These include using phage display to select large libraries of TCRs with randomized amino acids at specific positions are panned against NY-ESO-1 ^41–46^. Alternatively, in-silico protein engineering methods have also been used ^47^. Due to specific amino acid substitutions within the CDR1 and CDR2 regions of the α and β chain, these TCRs can possess significantly higher affinities than naturally occurring TCRs, reaching the picomolar range ^42,43^. However, such TCRs carry an inherent risk of cross-reactivity, potentially targeting peptides displayed on MHCs other than the intended epitope ^44,48–51^. Additionally, high affinity can induce T cell dysfunction ^52,53^, and reactivity towards features of the MHC molecules in the absence of cognate peptides ^54–56^.

Machine learning predictors can help identify epitope-specific TCRs within vast pools of potential candidates, such as TCR repertoires. TCR-epitope interaction predictors range from distance-based classifiers ^13,57,58^ to machine learning or deep learning models ^10,24,59–67^. TCR-epitope interaction predictor tools have been shown to identify epitope-specific TCRs with good accuracy if a large number of TCRs are available for a given epitope (approximately 50-100 TCRs) ^10,68^ but struggle to achieve robust predictions for epitopes for which TCR data is scarce or absent ^16,67,69,70^. For these reasons, as of today, robust predictions can only be performed for a few dozens of epitopes ^16,71^.

In this study, we designed a phage display experiment to collect a large number of TCRs recognizing the NY-ESO-1 epitope. Integrating this data into a machine learning TCR-epitope interaction predictor enabled us to identify epitope-specific TCRs showing activity towards NY-ESO-1 and no detectable cross-reactivity directly from TCR repertoires.

## Results

### Phage display reveals CDR3β binding motifs of TCRs specific for NY-ESO-1

To decipher the specificity of TCRs recognizing the NY-ESO-1 epitope, we built large phage display libraries of TCRs with randomized CDR3β loops. As a template, we first used a naturally occurring TCR targeting the NY-ESO-1 epitope, known as 1G4, which was isolated from the TCR repertoire of a melanoma patient ^32^. We further included a second and third template consisting of two affinity-enhanced TCRs, namely the 1G4-c50 and 1G4-c53c50 TCRs (Figure 1A and Table 1) ^43^. These two TCRs are characterized by amino acid substitutions in the CDR2 regions of 1G4 which interact with the MHC and significantly enhance the TCR affinity towards NY-ESO-1 ^43^ (Figure 1A).

**Figure 1.**
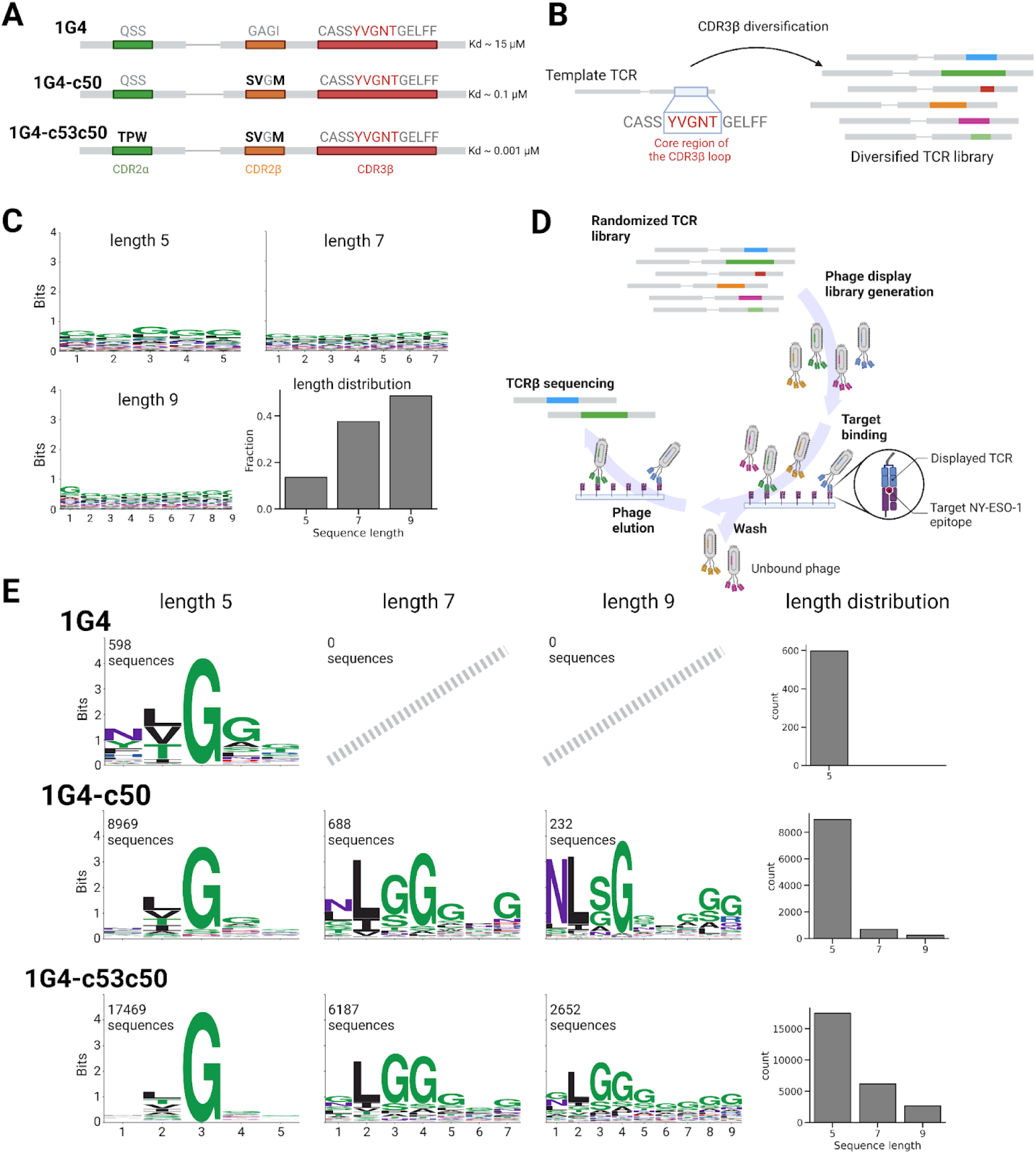
Phage display reveals CDR3β binding motifs of TCRs specific for NY-ESO-1. (A) Description of the three template TCRs used in the phage libraries. Amino acid substitutions of the 1G4-c50 and 1G4-c53c50 templates are highlighted in bold. The core region of the CDR3β (YVGNT) is highlighted in red. (B) Schematic of the design of the randomized TCR libraries for the phage display experiments. The TCRs have random amino acid sequences of length 5,7 and 9 in the core region of the CDR3β loops. (C) Sequence motifs and length distribution of the core region of CDR3β loops in phage libraries. (D) Illustration of the phage display experiment. The randomized TCRs expressed in phages were panned against the NY-ESO-1 pMHC monomer and sequenced. (E) Motifs and length distributions of the core regions of the CDR3β loops resulting after selection of phage libraries and motif deconvolution.

**Table 1.** Sequences of the 1G4, 1G4-c50, and 1G4-c53c50 template TCRs.

|  | Alpha chain [TRAV21*01/TRAJ6*01] |  |  | Beta chain [TRBV6-5*01/TRBJ2-2*01] |  |  |
| --- | --- | --- | --- | --- | --- | --- |
| Template TCR | CDR1 | CDR2 | CDR3 | CDR1 | CDR2 | CDR3 |
| 1G4 | DSAIYN | IQSSQRE | CAVRPTSGGSYIPTF | MNHEY | SVGAGI | CASSYVGNTGELFF |
| 1G4-c50 | DSAIYN | IQSSQRE | CAVRPTSGGSYIPTF | MNHEY | SVSVGM | CASSYVGNTGELFF |
| 1G4-c53c50 | DSAIYN | ITPWQRE | CAVRPTSGGSYIPTF | MNHEY | SVSVGM | CASSYVGNTGELFF |

For each template TCR separately, the core region of the CDR3β loop (corresponding to the YVGNT 5 amino acid sequence in 1G4, which are known to directly interact with the NY-ESO-1 epitope) was diversified with a two-step PCR process (Figure 1B and Supplementary Figure 1). The initial PCRs used a common forward primer and discrete reverse primers incorporating tails comprising different lengths of diversified trimer-defined codons. The chosen amino acid composition was designed to reflect that of naturally-occurring TCRs (Supplementary Table 1). These individual length-variant PCR products were then used in a second PCR reaction to introduce a XhoI restriction site downstream of the diversified CDR3β to facilitate substitution cloning into the TCR-containing vector (see *Methods,* Supplementary Figure 1 and Supplementary Table 2).

The randomized TCR library has a theoretical diversity larger than 10^8^ for each template, with randomized regions in the CDR3β loops of length 5, 7, and 9 (see *Methods*). To assess the quality of our input phage libraries, we sequenced them before any selection step (see *Methods* and Supplementary Data 1). Figure 1C shows the sequence motifs and the length distribution of the randomized regions merging the data for the three template TCRs. The N-terminal part (CASS) and C-terminal part (GELFF) of the CDR3β were not randomized since they do not directly interact with the epitope.

The TCRs libraries were incorporated into phages and panned against the NY-ESO-1 pMHC monomer immobilized on magnetic beads (Figure 1D). One round of panning was performed, incorporating different stringencies controlled by varying the number of wash cycles (1, 3 and 5 washes) (see *Methods*). The enrichment of specific CDR3β sequences was assessed by sequencing the panned phage libraries after each wash cycle (see *Methods* and Supplementary Figure 2).

CDR3β sequences obtained with 1, 3 and 5 washes were merged together as no significant differences were observed by varying the number of wash cycles (see *Methods* and Supplementary Figure 2). A significant number of unspecific TCRs are expected after panning and washing the phage libraries. To filter out these putative contaminants we used motif deconvolution with MoDec ^72^ (see *Methods*, Supplementary Figure 3 and Supplementary Data 2). The final binding motifs are reported in Figure 1E separately for each template and each length of the core region of the CDR3β loops. With the 1G4 template we already obtained several NY-ESO-1 specific TCRs (598 unique sequences). The 1G4-c50 and 1G4-c53c50 templates yielded a much higher number (9,889 and 26,308 unique sequences respectively). This aligns with expectations from their different intrinsic affinities (Figure 1A). The sequence motifs displayed high similarities across all templates and lengths, with enrichment of hydrophobic amino acids (Leu, Ile, Val) at position 2 and Gly at position 3 (Figure 1E). With the 1G4 template, only 5-mers could be retrieved in our pipeline (Figure 1E). On the contrary we obtained binding sequences for all three lengths with the 1G4-c50 and 1G4-c53c50 templates, albeit with a significantly higher number of 5-mers.

Overall, this analysis reveals that reproducible CDR3β motifs can be obtained by expressing large libraries of TCRs with randomized CDR3β loops in phage, and panning them with the NY-ESO-1 epitope.

### Integrating phage display data with machine learning tools enables robust predictions of NY-ESO-1 specific TCRs

We leveraged the TCRβ sequences obtained in the phage display to train a TCR-epitope interaction predictor for NY-ESO-1. To this end, we used the MixTCRpred machine learning framework ^10^ and trained a specific model for this epitope (see *Methods* and Figure 2A). As positives, we used all unique NY-ESO-1 specific CDR3β obtained with the 1G4, 1G4-c50, and 1G4-c53c50 templates. As negatives, we used CDR3β sequences from the randomized TCR libraries that did not bind NY-ESO-1 in the phage display experiment (see *Methods*). For quality control, we first performed a standard 5-fold cross-validation. As expected from the highly specific motifs in Figure 1E, we obtained high Area Under the receiver operating Curve (AUC) values, with a mean AUC of 0.97 (Figure 2B).

**Figure 2.**
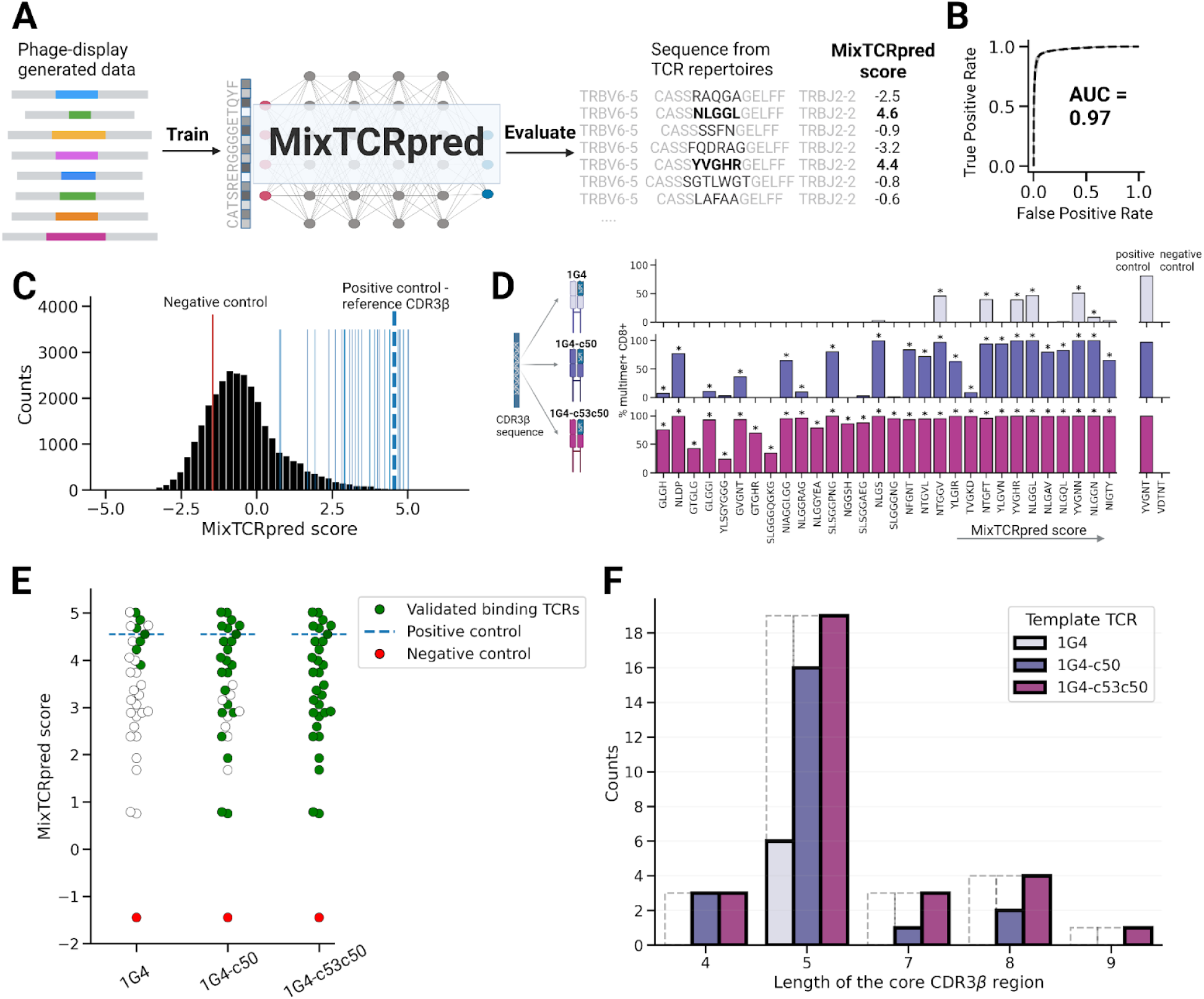
Integrating phage display data with machine learning tools enables robust predictions of NY-ESO-1 specific TCRs. (A) Illustration of training of MixTCRpred with CDR3β sequences obtained with the phage display screening, and evaluation of sequences from TCR repertoires of donors. (B) ROC curves obtained with a 5-fold cross-validation based on the phage display data. The dashed black line is the mean ROC curve. (C) Distribution of the MixTCRpred scores of CDR3β sequences from TCR repertoires. The blue lines show the scores of the 30 CDR3β sequences selected for experimental testing. The dashed blue line shows the score of the reference CDR3β sequence CASS**YVGNT**GELFF. (D) Percentage of multimer+CD8+ cells among Jurkat cells transduced with each of the 30 TCRs selected in panel C. TCRs are labeled based on the sequence of the core region of CDR3β loops, and ordered by the MixTCRpred scores. Stars indicate TCRs considered as NY-ESO-1 specific. (E) MixTCRpred scores of the TCRs with different CDR3β loops that could (green) or could not (white) be experimentally validated with the 1G4, 1G4-c50, and 1G4-c53c50 templates. The MixTCRpred scores of the positive control (the reference CDR3β sequence CASS**YVGNT**GELFF) and of the negative control (the CASS**VDTNT**GELFF sequence) are also shown. (F) Length of the core region of the CDR3β sequences that were tested (dashed lines) and validated (solid lines) for the three TCR templates.

We next explored whether our MixTCRpred model could be used to identify NY-ESO-1 specific TCRs directly from TCR repertoires. To this end, we first collected a large number of TCRβ sequences from TCR repertoires of unrelated donors ^73^. To be consistent with the design of our phage display libraries, we only included TCRβ with TRBV6-5 and TRBJ2-2 genes (see *Methods*). In total, we retrieved 29,867 TCRβ sequences, which were scored with our MixTCRpred model. The distribution of the MixTCRpred scores is shown in Figure 2C. TCRβ with high scores are predicted to be NY-ESO-1 specific. The reference CDR3β sequence (CASS**YVGNT**GELFF) has a MixTCRpred score of 4.55, ranking among the top-scoring sequences (Figure 2C).

To investigate and experimentally validate the potential of our *in-silico* predictions, we selected 30 TCRβ with a broad range of MixTCRpred scores and including different lengths for the core region of the CDR3β loop (Figure 2C and Table 2). We also included the reference CDR3β sequence (CASS**YVGNT**GELFF) as positive control, and a randomly selected CDR3β (CASS**VDTNT**GELFF) having a score of -1.45 to use as negative control. All TCRs with each of the three templates (i.e., 1G4, 1G4-c50, and 1G4-c53c50) were tested for binding to the NY-ESO-1 epitope (SLLMWITQC) (Figure 2D). To this end, RNA encoding each of the selected TCRs was synthesized and introduced into Jurkat cells via electroporation. Following overnight incubation, the TCR-transfected cells were interrogated for binding with NY-ESO-1-multimers (see *Methods* and Figure 2D). An illustration of the results of the multimer staining is shown in Supplementary Figure 4. A relatively small number of TCRs could be validated with the 1G4 template (6 out of 30) with variable percentages of multimer+CD8+ Jurkar cells from the multimer staining experiment (Figure 2D). All validated TCRs ranked among the top-scoring predictions with MixTCRpred (Figure 2E), and had core regions in the CDR3β loop of length 5 (Figure 2F). Conversely, most (i.e., 22 out of 30) TCRs with the 1G4-c50 template and all TCRs with the 1G4-c53c50 template were found to bind to NY-ESO-1 (Figure 2D-E). TCRs with CDR3β of multiple lengths could be validated with the affinity-enhanced templates (1G4-c50 and 1G4-c53c50), including some with lengths not included in the training set of our MixTCRpred model (Figure 2F).

**Table 2.**
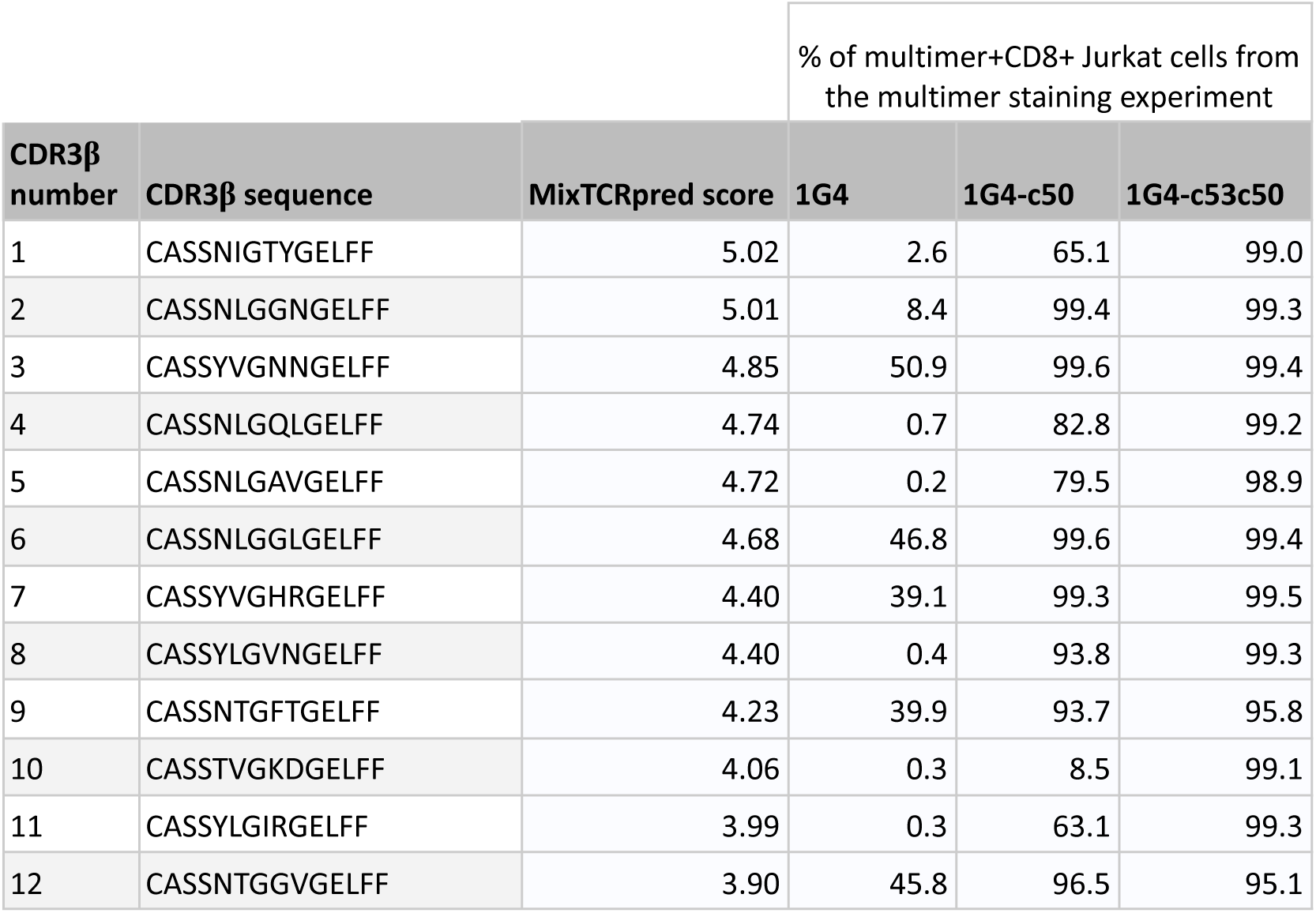
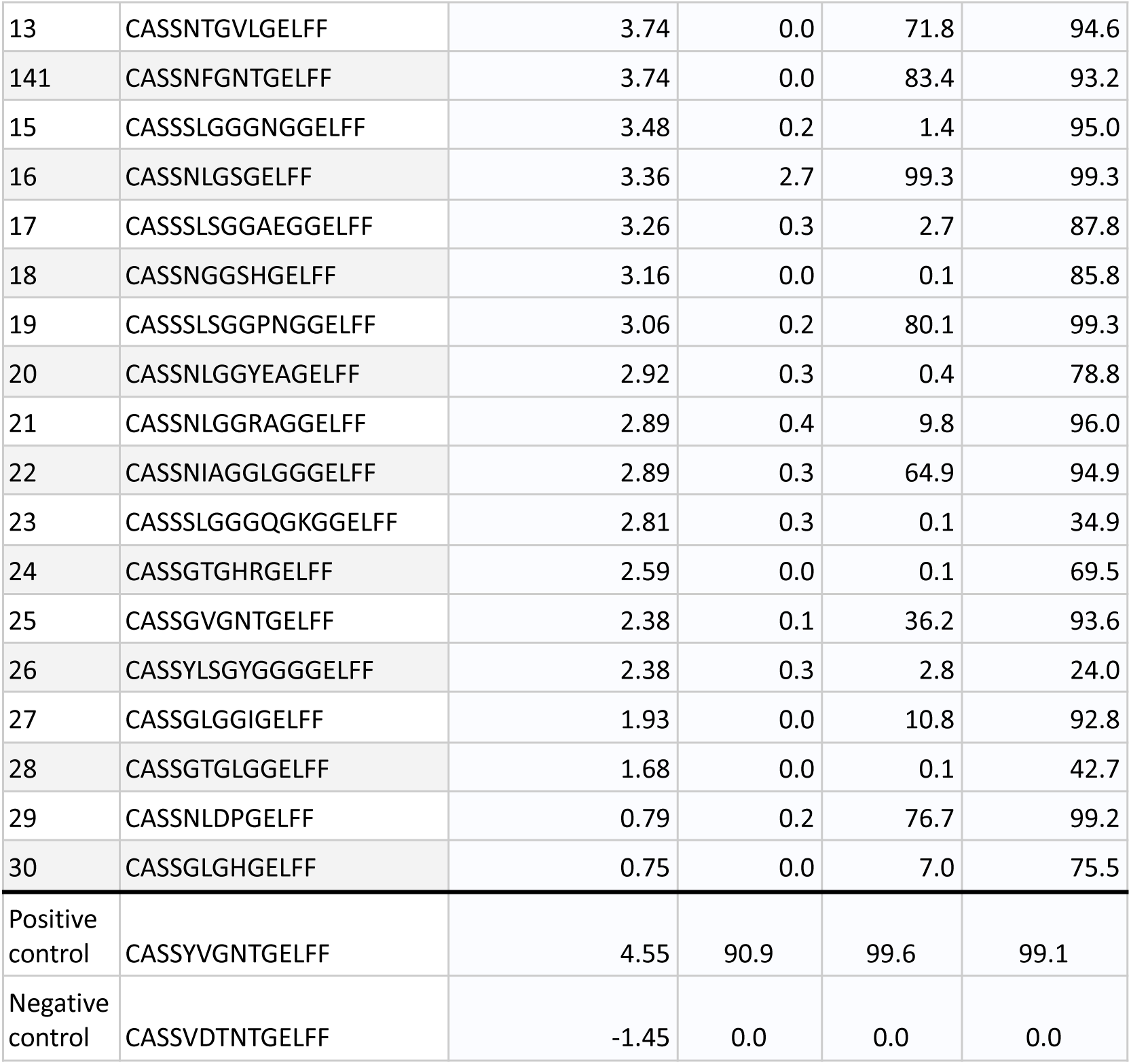
List of CDR3β selected for experimental validation together with the MixTCRpred scores and the percentages of multimer+CD8+ T cells resulting from the multimer staining experiments with the three template TCRs, averaged across two repetitions.

To assess how using different templates in the phage libraries influenced the predictive power of MixTCRpred, we trained three template-specific models. Each model used as positives TCR sequence data obtained from the phage display screening with a specific template (see *Methods* and Supplementary Figure 5). Despite the highly variable number of training data (598 positive with the 1G4 templates, 9889 with 1G4-c50, and 26,308 with 1G4-c53c50) the TCRs validated for NY-ESO-1 binding consistently ranked as top-scoring sequences for each of the three models (Supplementary Figure 5).

This analysis demonstrates that the data obtained with the phage display pipeline can be effectively used to train a predictive model, which can then be used to identify NY-ESO-1 specific TCRβ sequences directly from TCR repertoires.

### MixTCRpred trained on phage display data outperforms other approaches for predictions of NY-ESO-1 specific TCRs

We next compared our strategy for identifying NY-ESO-1 specific TCRs with other approaches. To this end, we capitalized on the 30 experimentally tested TCRs in Figure 2D with the 1G4 template (i.e., 6 positives and 24 negatives) and on the fact that they span a large range of MixTCRpred scores (i.e., were not restricted to the top scoring TCRs). MixTCRpred trained on the phage display data generated with the three template TCRs (29,688 NY-ESO-1 specific TCRs) achieved an AUC of 0.88 (Figure 3A). Using as training set only phage display data obtained with the naturally occuring 1G4 template (598 NY-ESO-1 specific TCRs) yielded an AUC of 0.92 (Figure 3A). Another method consists of assessing which of the 30 tested CDR3β has an exact match in the TCRs observed in the phage display experiments. We obtained an AUC of 0.58 with this approach (Figure 3B). As an alternative to using the results of the phage display experiment, we calculated the sequence similarity of the 30 TCRs with the reference CDR3β (CASS**YVGNT**GELFF) using the TCRbase and tcrdist3 distance metrics ^57,74^. Some validated CDR3β sequences have high sequence similarity to the reference CDR3β, while others have lower sequence similarity. Overall, we obtained an AUC of 0.79 for TCRbase and of 0.68 for tcrdist3 (Figure 3C). As a final test, we investigated whether existing machine learning predictors could have identified the validated NY-ESO-1 specific TCRs. We used three predictors (NetTCR2.2 ^68^, epiTCR ^75^ and pMTNet ^68,76^) which include in their training set the few publicly available TCRs known to bind to NY-ESO-1. All three achieved AUCs lower than 0.7 (Figure 3D).

**Figure 3.**
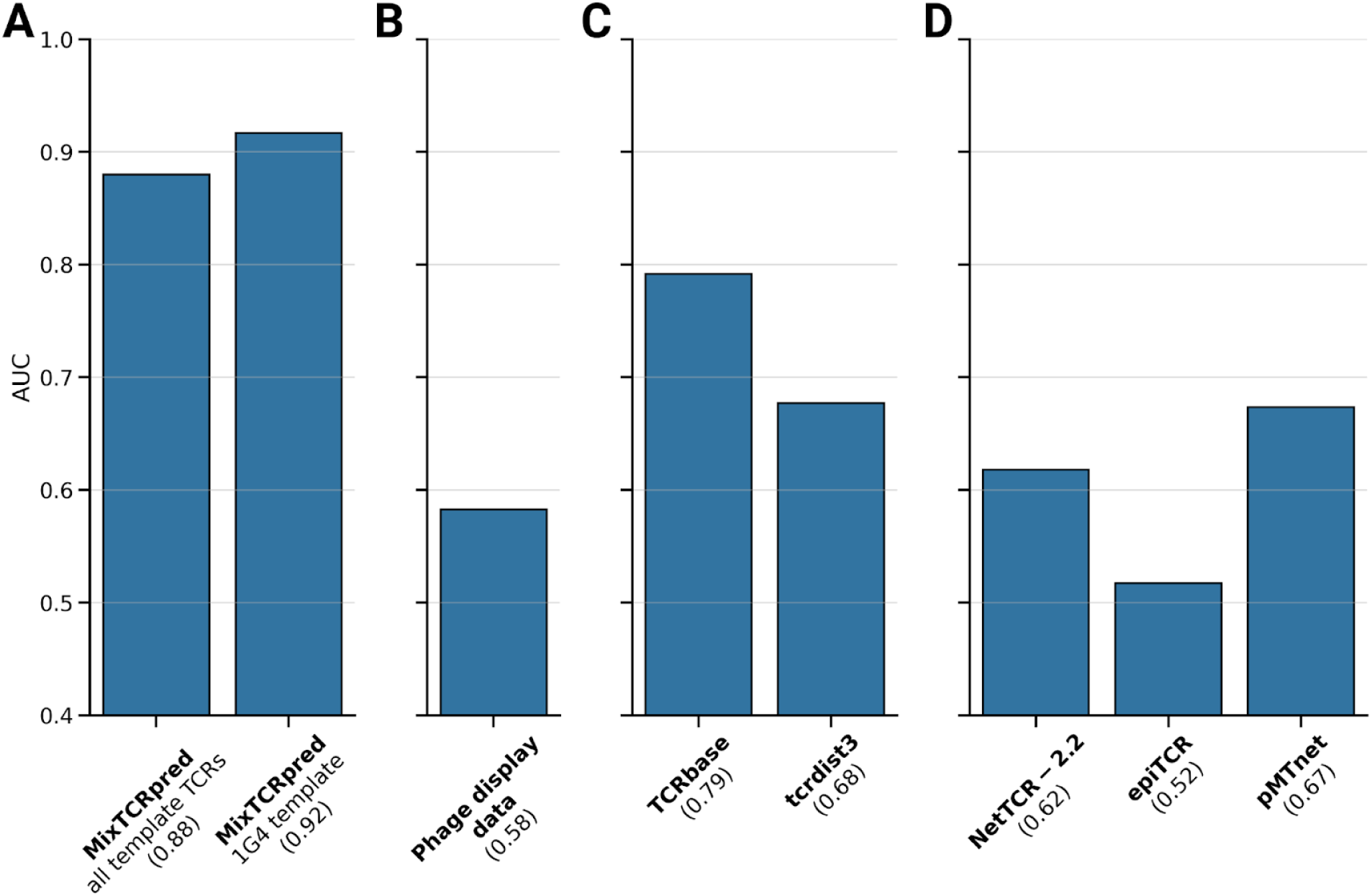
MixTCRpred trained on phage display data outperforms other approaches for predictions of NY-ESO-1 specific TCRs. (A) AUC achieved by MixTCRpred trained on the phage display data generated with the three template TCRs (1G4, 1G4-c50 and 1G4-c53c50) and with the 1G4 template exclusively. (B) AUC obtained by looking for an exact match of the TCRs in the data generated by phage display. (C) AUCs achieved by computing sequence similarity to the reference CDR3β (CASS**YVGNT**GELFF) with TCRbase or tcrdist3. (D) AUCs obtained with three pre-trained pan-epitope predictors which do not include the data generated with the phage display in their training sets.

Overall, this benchmark shows that combining phage display data with machine learning represents a promising strategy for identifying TCRs binding to NY-ESO-1 within a pool of potential candidates.

### TCRs identified by MixTCRpred display activity towards NY-ESO-1 and no detectable cross-reactivity

To investigate the functionality of the TCRs predicted by MixTCRpred to bind to NY-ESO-1 we conducted multiple cellular activation assays. To this end, we selected three predicted TCRs (CDR3β: CASSYVGN**N**GELFF, CASSYVG**HR**GELFF, CASS**NL**G**GL**GELFF, see Table 2) as well as the template 1G4 (CDR3β: CASSYVGNTGELFF). All these TCRs were among the top MixTCRpred predictions and were validated as NY-ESO-1 binders on the 1G4 template.

We synthesized DNA encoding for these TCRs and transduced them into Jurkat cells. Jurkat cells were co-cultured overnight with HLA-A*02:01 positive T2 cells pulsed with the NY-ESO-1 peptide. Activation markers CD69 and PD-1 were used to identify peptide-activated T cells. This experiment showed that Jurkat cells expressing any of the four TCRs were specifically activated by the NY-ESO-1 epitope (Figure 4A). On the contrary, these cells were minimally activated (close to zero fraction of CD69^+^PD-1^+^ T cells) by the CMV-derived epitope NLVPMVATV which was used as a negative control.

**Figure 4.**
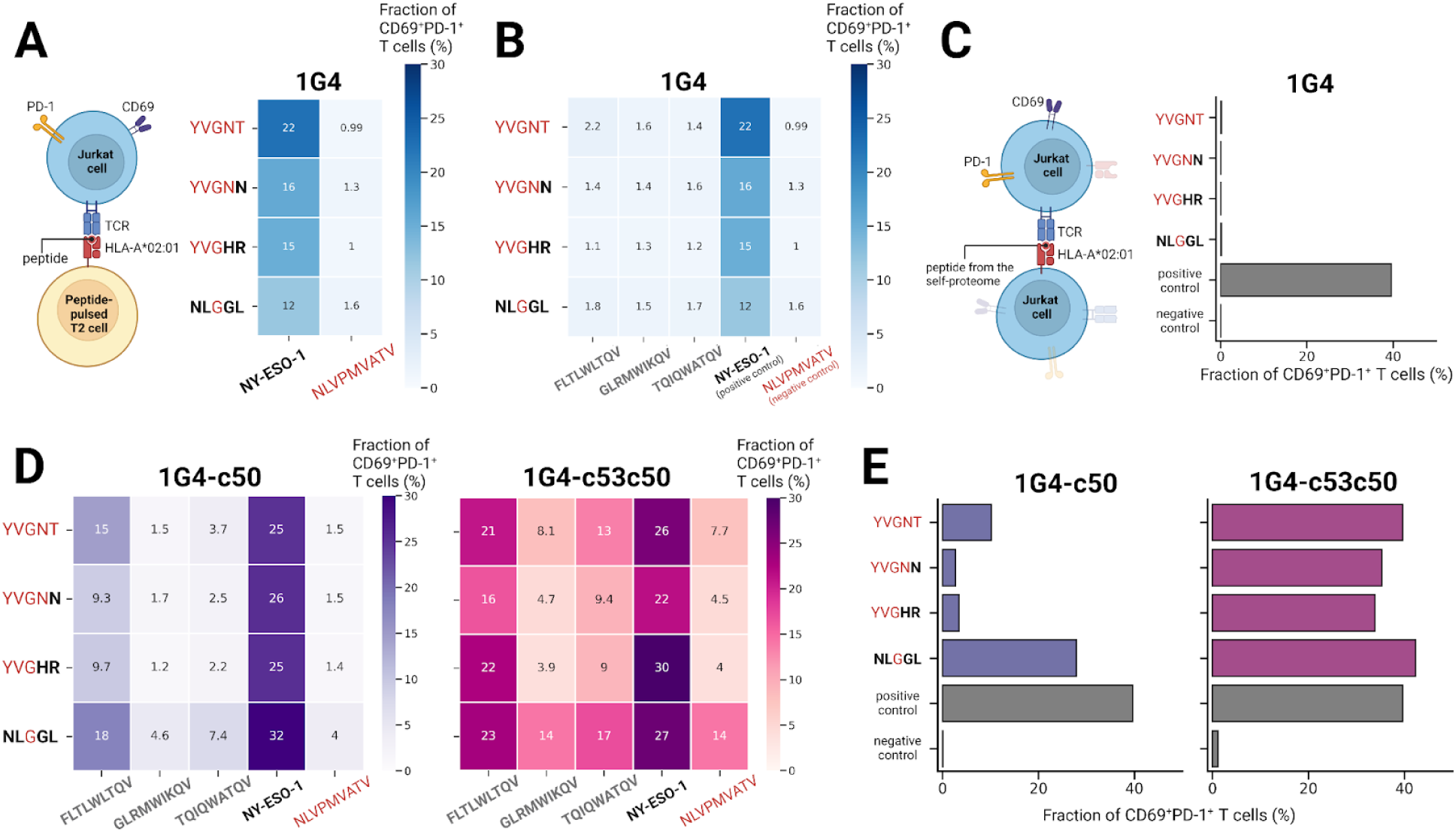
TCRs identified by MixTCRpred display activity towards NY-ESO-1 and no detectable cross-reactivity. (A) Heatmap showing the fraction of CD69+PD-1+ Jurkat cells encoding four TCRs with different CDR3β sequences based on the 1G4 template after co-culture with T2 cells pulsed with the NY-ESO-1 epitope at physiologically relevant peptide concentration (0.1 μg/mL). As negative control, the results for the CMV-derived epitope NLVPMVATV are also shown. For clarity, only the sequence of the core region of the CDR3β loop is shown. (B) Heatmap showing the fraction of CD69+PD-1+ Jurkat cells encoding the four TCRs of panel A after co-culture with T2 cells pulsed with three peptides from the self proteome at peptide concentration 0.1 μg/mL. The NY-ESO-1 peptide was used as positive control, and the CMV-derived epitope NLVPMVATV as negative control. (C) Fraction of CD69+PD-1+ Jurkat cells expressing the four TCRs of panel A and activated by co-culturing with HLA-A*02:01 positive Jurkat cells presenting peptides derived from the self-proteome. Jurkat cells without any TCR expression (no transduction) were used as negative controls. Jurkat cells expressing the high-affinity TCR 1G4-c53c50 were used as positive control. (D) Heatmap showing the fraction of CD69+PD-1+ Jurkat cells encoding four TCRs based on the affinity-enhanced 1G4-c50 and 1G4-c53c50 templates when stimulated by the indicated peptide at peptide concentration 0.1 μg/mL. (E) Fraction of activated Jurkat cells encoding four CDR3β sequences based on the affinity-enhanced 1G4-c50 and 1G4-c53c50 templates when stimulated by peptides derived from the self-proteome.

To assess putative cross-reactivity, we investigated whether our TCR-transduced Jurkat cells could be activated by self-peptides displaying similarity to NY-ESO-1. We first selected three peptides (FLTLWLTQV, GLRMWIKQV, and TQIQWATQV) from the human proteome predicted to be presented by HLA-A*02:01, and having high sequence similarity with the NY-ESO-1 epitope (SLLMWITQC, see *Methods*). Two of them (FLTLWLTQV, TQIQWATQV) were also reported in an earlier study investigating the binding properties of the affinity-enhanced TCR NY-ESOc259 ^37^. The peptides were pulsed on HLA-A*02:01 positive T2 cells and incubated overnight with Jurkat cells transduced individually with the three predicted TCRs (i.e., CDR3β loops CASSYVGN**N**GELFF, CASSYVG**HR**GELFF, CASS**NL**G**GL**GELFF on the 1G4 template). We observed specific activation by the NY-ESO-1 epitope and close to zero activation by the other three peptides. The residual cross-reactivity was even lower than for the template TCR 1G4 with the reference CDR3β (CASSYVGNTGELFF) (Figure 4B). These results were confirmed at various peptide concentrations (Supplementary Figure 6).

To broaden our cross-reactivity investigation beyond the few selected peptides, we performed a functional assay to evaluate T cell cross-reactivity towards peptides from the self-proteome. Jurkat cells were dually transduced for both TCRs and HLA-A*02:01 and maintained under steady-state culture conditions for 3 to 6 days. Such cells spontaneously present epitopes derived from a multitude of endogenously expressed proteins, which could induce T cell activation through the transduced TCR (Figure 4C). Jurkat cells without any TCR expression (no transduction) were used as negative controls while cells expressing the high-affinity TCR 1G4-c53c50 were used as positive control (see *Methods*). The three predicted TCRs mirrored the template 1G4 TCR in triggering minimal T cell activation in this assay (close to zero fraction of CD69^+^PD-1^+^ T cells), suggesting a strong retention of specificity for the cognate target peptide (Figure 4C).

We next repeated these experiments with the three predicted TCRs, and the one with the reference CDR3β, this time based on the affinity-enhanced templates 1G4-c50 and 1G4-c53c50. We observed that TCR-transduced Jurkat cells were functionally activated by the NY-ESO-1 peptide, but displayed cross-reactivity with the FLTLWLTQV and, to a lesser extent, with the GLRMWIKQV and TQIQWATQV peptides (Figure 4D and Supplementary Figure 6). Moreover, these TCR-Jurkat cells showed cross-reactivity towards HLA-A*02:01-presented peptides of the Jurkat self-proteome (Figure 4E).

Overall, this analysis reveals that TCRs predicted by MixTCRpred exhibit activity towards NY-ESO-1 and no detectable cross-reactivity. Reversely, cross-reactivity with both specific peptides and the peptidome was detected when using Jurkat cells transduced with the affinity-enhanced 1G4-c50 and 1G4-c53c50 TCR templates.

### Structural analyses reveal the molecular basis of the CDR3β binding motifs

To gain molecular insights into the binding motifs of NY-ESO-1 specific TCRs, we first attempted to model the three TCRs functionally validated in Figure 4A (CDR3β: CASSYVGN**N**GELFF, CASSYVG**HR**GELFF and CASS**NL**G**GL**GELFF) using as template the crystal structure of 1G4 (PDB: 2BNR). Overall, the CDR3β loops are predicted to maintain a conformation similar to that of 1G4 (CDR3β: CASSYVGNTGELFF). Even in the case of four point mutations (**NL**G**GL** versus YVGNT), the mutations Y94N and V95L in the first two residues did not significantly alter the positions of the α-carbons with respect to the reference CDR3β. Slightly more significant structural rearrangements are predicted to occur towards the end of the core regions of the CDR3β loop (Figure 5A), which is consistent with the lower amino acid specificity at these positions in the motifs of Figure 1E. We next analyzed the beta-factors of Cα across the CDR3β loop (Figure 5B). Higher beta-factors were observed for the last 2 amino acids, which is consistent with the lower specificity and higher flexibility observed in the motif and structural models of CDR3β loops with a core region of length 5.

**Figure 5.**
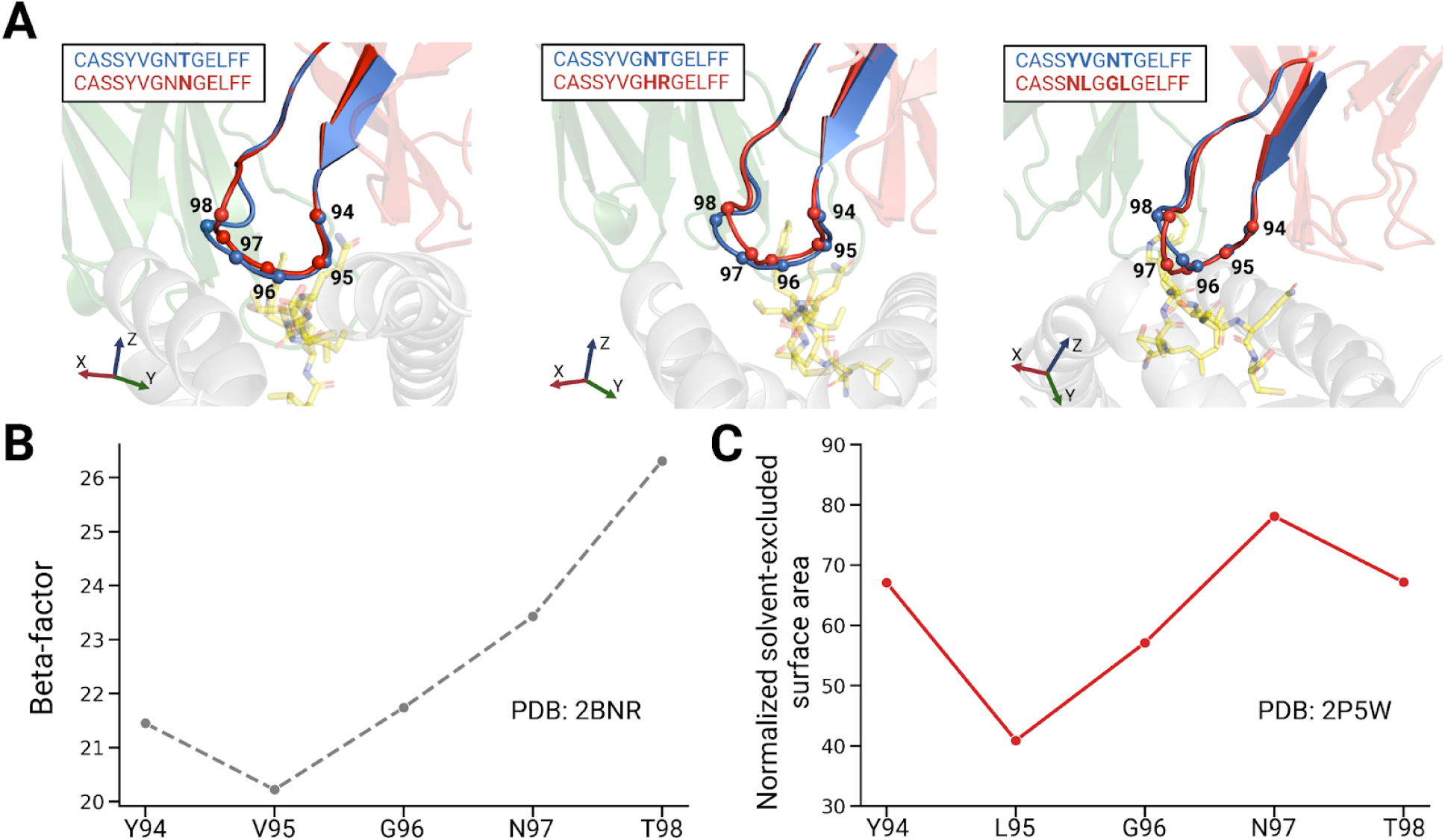
Structural analyses reveal the molecular basis of the CDR3β binding motifs. (A) Predicted binding modes of three CDR3β loops (CASSYVGNNGELFF, CASSYVGHRGELFF and CASSNLGGLGELFF, in red), overlapped with the reference CDR3β (CASSYVGNTGELFF, in blue). The molecular modeling was performed using the crystal structure of the 1G4 template (PDB: 2BNR). The α-carbons of the core region of the CDR3β loops are represented as spheres. (B) Beta factor of the core region of the CDR3β loop (PDB: 2BNR). (C) Normalized solvent-excluded surface area of amino acid side chains in the core region of CDR3β loop in the 1G4-c53c50 template (PDB: 2P5W, residue numbering based on PDB: 2BNR).

We next explored potential mechanisms for the longer CDR3β loops observed in the CDR3β binding motifs based on affinity-enhanced templates (Figure 1E) and in some of the TCRs validated on these templates (Figure 2D). To determine where the CDR3β loops may accommodate the additional residues, we analyzed the solvent accessibility of amino acid side chains in the core region of the CDR3β loop for the 1G4-c53c50 template (PDB: 2P5W). In this crystal structure Leu is found at the second position (L95) instead of Val, which is compatible with the sequence motifs we identified in the phage display experiments (Figure 1E). We observed that amino acids located in the C-terminal region of the core of the CDR3β loop (position 97 and 98) are more solvent-exposed (Figure 5C). This suggests that the loop extension occurs towards the end of the core region of the CDR3β loop. This is consistent with the longer motifs for the affinity-enhanced templates which were obtained in the phage display experiments (Figure 1E).

Overall, our structural analyses provide a molecular interpretation for the high specificity and conservation across lengths of the N-terminal part of the binding motifs in Figure 1E and the lower specificity at the C-terminal part.

## Discussion

Phage display provides a powerful framework to screen very large libraries of TCRs with randomized amino acids at specific positions against clinically relevant epitopes. In this work, we demonstrate that phage display can be used in combination with machine learning predictors of TCR-epitope interactions to identify epitope-specific TCRs directly from TCR repertoires.

Our phage display approach relies on having at least one TCR specific for the epitope under investigation, referred to as the template TCR, from which a library with randomly mutated CDR3β loops can be designed and selected against a specific epitope. Analysis of epitope-specific TCRs in VDJdb shows that this is the case for more than 1000 epitopes. The phage display approach holds particular significance for cancer epitopes which are typically challenging to experimentally identify in primary T cells from patients or healthy donors. These include the cancer-testis antigen NY-ESO-1_157–165_.

Sequencing data from high-throughput phage display screening can contain a large number of putative contaminants. Our results show that motif deconvolution algorithms like MoDec ^72^ can be effectively used to unravel binding motifs even in the presence of a substantial fraction of unspecific TCRs. The high similarity of the motifs observed in all three templates as well as our multiple experimental validations demonstrate that motifs identified by motif deconvolution in this work represent *bona fide* CDR3β binding motifs of TCRs specific for the NY-ESO-1 epitope.

Earlier studies ^42,43^ used iterative selection and amplification of phage libraries to identify TCRs with enhanced affinity for different epitopes by exploring amino acid substitutions at multiple positions across the TCR-epitope interaction interface (i.e., not restricted to the variable region of the CDR3 loops). As a result, many of the V segments of these TCRs deviate from those of native TCRs and have not undergone positive and negative selection in the thymus. This increases the risk of cross-reactivity with self-peptides, as demonstrated in this work for the affinity-enhanced 1G4-c50 and 1G4-c53c50 templates, potentially leading to toxicity in clinical applications ^49^. Naturally-occurring TCRs from TCR repertoires represent promising candidates for the development of T-cell-based immunotherapies ^77^, having a lower risk of cross-reactivity and thus a reduced risk of adverse reactions compared to affinity-enhanced TCRs. Our proposed strategy - consisting of designing phage display libraries specific for the most variable region of the CDR3β loop and using the data for training TCR-epitope interaction prediction tools to interrogate TCR repertoires - enabled us to rapidly identify multiple candidates within tens of thousands of TCR sequences. As such, the machine learning MixTCRpred model could facilitate and accelerate the identification of multiple TCRs for adoptive transfer in cancer immunotherapy. Although we cannot rule out potential cross-reactivity with peptides presented in specific human tissues and not in our system based on HLA-A*02:01 expressing Jurkat cells, our results already enable us to exclude cross-reactivity with a significant fraction of the human HLA-A*02:01 self-peptidome.

Our study shows that high accuracy in predicting NY-ESO-1 specific TCRs can be achieved by combining our high-throughput experimental phage display pipeline together with a customized machine learning model. Other approaches which either do not use the phage display data or do not have a dedicated machine learning model achieved lower accuracies, as demonstrated in our internal benchmark. For this benchmark, we used experimentally tested TCRs with a broad range of MixTCRpred scores, thereby mitigating biases towards the high-score predictions. We cannot exclude that some high-scoring TCRs with other tools and with CDR3β sequences incompatible with the phage-derived motifs may still bind but were not included in our test set. However, the depth of the phage display libraries suggest that these cases should be rare.

The binding motifs derived from the phage display experiments with different templates displayed high similarity. This observation has several important consequences. First, it shows that affinity-enhanced templates are not necessary for the pipeline proposed in this work. As such, many native TCRs binding to clinically relevant epitopes in public databases could be used as templates for designing phage display libraries similar to the one built in this work. Second, it shows that affinity-enhanced templates, which displayed extensive cross-reactivity and therefore would not be suitable for any clinical application, are compatible with the training of predictors and do not lead to artifacts when scoring TCRs from TCR repertoires. This may be useful when working with clinically relevant epitopes for which only very low affinity TCRs are known and which could not be used directly in phage display.

In our phage display libraries, we diversified the core region of the template CDR3β loop across different lengths, keeping the V and J genes as well as the CDR3α loop unmodified. As a result, the MixTCRpred model trained on such data is only applicable to make predictions for TCRβs with these specific V and J genes. While this represents a limitation of the current study, we anticipate that the proposed combined experimental and computational framework will be useful both for designing and screening phage display libraries built with other V and J genes and/or including mutated regions in the CDR3α loop.

In summary, our work presents an integrated experimental and computational pipeline to identify TCRs recognizing clinically relevant epitopes. In particular, we demonstrate that combining phage display with machine learning enabled us to train TCR-epitope interaction predictors for a clinically-relevant epitope for which only few TCRs had been identified by other means, and interrogate TCR repertoires. We anticipate that this work will pave the way for designing larger TCR libraries encoded in phage or other organisms ^79,80^ to expand the epitope coverage of TCR-epitope interaction predictors and facilitate the identification of epitope-specific TCRs directly from TCR repertoires of cancer patients or healthy donors.

## Methods

### Randomized CDR3β library

Randomized CDR3β libraries of DNA of differing lengths were constructed on the 1G4 single chain (sc)TCR scaffold. Briefly, 1G4 TCR domains were codon-optimized for expression in *E. coli* and cloned as a 3-domain Vα-VβCβ single-chain ORF fused N-terminally to the M13/f1 gIII major coat protein in a pUC19-based phagemid vector (pCHV101). In addition to parental 1G4, which has an affinity for its cognate antigen (NY-ESO-1_157-165_) of ∼10μM ^38–40^, we also included two variants containing CDR2 “anchoring” mutations which have been shown to increase the affinity of the TCR to ∼0.1μM and ∼0.001μM respectively through improved contacts with the common HLA-A*02:01 helices that flank the peptide groove ^43^. To circumvent the reported instability conferred by the Cys at 3’ of the NY-ESO-1 peptide, we conducted all phage library selection using the more stable A0201_SLLMWITQ**V** peptide-MHC complex ^81^. Using these scTCR DNA templates, the core region of the 1G4 CDR3β (YVGNT) was diversified in a two-step PCR using standard procedures. The initial PCRs used a common forward primer and discrete reverse primers incorporating tails comprising different lengths of trimer-defined (TRIM) codons (Ella Biotech GmbH, Martinsreid, Germany). We interrogated the publicly accessible VDJ database ^27^ comprising some 40K human TCRs targeting ∼1000 distinct MHC antigens. From this data, a single amino acid codon mixture was devised to approximate the composition of natural CDR3β loop cores (Supplementary Table 1). These individual length-variant PCR products were then used as templates in a second PCR reaction to introduce a XhoI restriction site downstream of the diversified CDR3 (Supplementary Figure 1 and Supplementary Table 2).

### Phage library

The diversified library PCR fragments were double digested with AscI (upstream of CDR3β) and XhoI and ligated into the similarly digested pCHV101 template vectors. Ligated products were purified and used to electroporate electrocompetent *E. coli* TG1 cells (Lucigen) which were then directly plated onto large 2TY-agar plates supplemented with 100 µg/ml ampicillin/2% glucose and incubated at 30 °C overnight. The following day, the bacterial libraries (all of size >10^8^ clones) were harvested from plates and stored as concentrated glycerol stocks at -80 °C.

Filamentous phage libraries were rescued using M13 helper phage (Life Technologies), concentrated and purified according to standard procedures and stored in aliquots at -80 °C. The physical size of the constructed libraries actually realized (number of bacterial clones on plates) were 2.7x10^8^; 2.6x10^8^; 6.1x10^8^ for 5-mers, 7-mers and 9-mers respectively. To assess diversity, the libraries were sequenced with 2x250bp paired-end NGS sequencing using an in-house Illumina MiSeq platform, requesting one million reads per library (Supplementary Data 1). 37.5%; 69% and 49.8% of clones in each library were determined to be functional intact clone opening reading frames identical to the parental TCR.

### Phage panning

Phage panning of the libraries was performed according to standard procedures against monomeric biotinylated HLA-A*02:01,NY-ESO_157–165_ (heteroclitic peptide variant: SLLMWITQV) immobilized on M-280 streptavidin magnetic beads (Thermo Fisher Scientific), using an irrelevant bio-HLA-A*0201 peptide complex for deselection of non-specific binders. One round of panning was conducted, incorporating different stringencies controlled by varying the number of wash cycles. The enrichment of specific CDR3β sequences was assessed by performing 2x250bp paired-end NGS sequencing on the extracted and amplified output DNA using an in-house Illumina MiSeq platform and requesting one million reads in each sample.

### Phage display data processing

The TCR-sequencing data obtained from the Illumina MiSeq platform were processed with the MiXCR v3.0.13 with standard parameters (mixcr analyze amplicon --species hs starting-material dna --5-end v-primers --3-end j-primers --receptor-type TRB) ^82^. To ensure high quality data for the phage display generated data, we removed all the TCR sequences occurring only once, which are likely due to sequencing errors. We also removed the “YVGNT” sequences from the phage display generated data, which could reflect template TCRs which failed randomization. Duplicated sequences were removed, retaining only unique TCR clonotypes. The unsupervised motif deconvolution tool MoDec was used to further process the TCR sequences ^72^. We set the number of motifs to 1 (k=1) and background frequencies based on the amino acid distribution in the input phage libraries. TCR sequences belonging to the flat motif or with motifs lacking positions with information content higher than 1, were considered as putative contaminants and removed (Supplementary Figure 2 and Supplementary Data 2).

### MixTCRpred - model architecture and data

We developed a MixTCRpred model (A0201_NY-ESO-1-CDR3b) customized for CDR3β sequence data used in this study. We used as positives the phage display output sequences from the 1G4, 1G4-c50, and 1G4-c53c50 templates (29,688 unique TCR sequences). As negatives (CDR3β non-binding to NY-ESO-1) we used 7,330 sequences from the phage input libraries that were not present in the output libraries. When training template-specific MixTCRpred models we used the data generated with each template separately (598 positive for the 1G4 templates, 9889 for 1G4-c50, and 26,308 for 1G4-c53c50). These data were used to train a new MixTCRpred model (A0201_NY-ESO-1-CDR3b), following the procedure previously described in ^10^, and the model accuracy was accessed with a standard 5-fold cross-validation. We collected a large number of TCRβ sequences from TCR repertoires downloaded from iReceptor ^73^with TRBV6-5 and TRBJ2-2 genes and filtered out sequences with N- and C-terminal parts different from that of the reference CDR3β (respectively CASS and GELFF) or containing non-standards amino-acids. In total we retrieved 29,867 TCRβ sequences.

### TCR cloning and multimer staining

30 CDR3β with high MixTCRpred scores were selected for pMHC multimer staining experiments (Table 2). CDR3β sequences were based on the three template TCRs (1G4, 1G4-c50, or 1G4-c53c50, Table 1), and tested for binding versus the NY-ESO-1 epitope (HLA-A*02:01,SLLMWITQC) (Figure 2D). TCRα/β pairs were cloned into Jurkat cells (TCR/CD3 stably transduced with human CD8α/β and TCRα/β CRISPR-KO) ^83,84^. Codon-optimized DNA sequences coding for paired α and β chains, were synthesized at GeneArt (Thermo Fisher Scientific) or elesis Bio DNA. The DNA fragments served as template for in vitro transcription (IVT) and polyadenylation of RNA molecules as per the manufacturer’s instructions (Thermo Fisher Scientific), followed by co-transfection into recipient T cells. Jurkat cells were electroporated using the Neon electroporation system (Thermo Fisher Scientific) with the following parameters: 1,325 V, 10 ms, three pulses. After overnight incubation, electroporated Jurkat cells were interrogated by pMHC-multimer staining with the following surface panel: anti-hCD3 APC Fire 50 (SK7, Biolegend Cat# 641415, 0.4μL in 50μL); anti-hCD8 FITC (SK-1 Biolegend, Cat# 344704, 0.15μL in 50μL); anti-hCD4 PE-CF594 (RPA-T4, BD Bioscience Cat# 562281, 0.4μL in 50μL); anti-mouse TCRβ-constant APC (H57-597, Thermo Fisher Scientific, Cat# 17-5961-81, 0.6μL in 50μL); pMHC-multimer-PE (HLA-A*02:01 with the SLLMWITQC peptide) in-house synthesized, 1μL in 50μL); viability dye Aqua (L34966, Thermo Fisher Scientific, 0.15μL in 50μL staining mix in PBS). The peptides and pMHC multimers were produced by the Peptide and Tetramer Core Facility of the Department of Oncology, UNIL-CHUV, Lausanne. Samples were acquired by flow cytometry and FACS data were analyzed with FlowJo 10.8.1 (TreeStar).

### Benchmarking MixTCRpred trained on phage display data with other approaches

We used the 30 experimentally tested TCRs with the 1G4 template (6 positives and 24 negatives) to benchmark our strategy with the following approaches:

- TCRbase 1.0 (web server: https://services.healthtech.dtu.dk/services/TCRbase-1.0/). The 1G4 template sequence (Table 1) was used as the training database and the list of the 30 tested TCRs were uploaded on the web server to compute the sequence similarity with the reference CDR3β (CASS**YVGNT**GELFF).
- Tcrdist3 (GitHub page: https://github.com/kmayerb/tcrdist3). The function compute_distances from the tcrdist package was used to compute the sequence similarity of the 30 tested TCRs with the reference CDR3β (CASS**YVGNT**GELFF).
- NetTCR2.2 (web server: https://services.healthtech.dtu.dk/services/NetTCR-2.2/). The web server was used to compute the likelihood of interaction between the NY-ESO-1 peptide and the 30 tested TCRs.
- epiTCR (GitHub page: https://github.com/ddiem-ri-4D/epiTCR). The pre-trained model models/rdforestWithoutMHCModel.pickle was used to compute the interaction score between the NY-ESO-1 peptide and the 30 tested TCRs.
- pMTNet (web server: https://dbai.biohpc.swmed.edu/pmtnet/). he 30 tested TCRs sequences were uploaded on the web server to compute the likelihood of interaction with the NY-ESO-1 peptide.

### Functional assay - cell lines and culture

TCR knock-out HLA-A*02:01^neg^/J76 CD8αβ cells (kindly provided by Drs. I. Edes and W. Uckert, Max-Delbrück-Center, Berlin, Germany), TCR knock-out HLA-A*02:01^pos^/J76 CD8αβ cells ^54^ and HLA-A*02:01^pos^ TAP-deficient T2 cells (ATCC CRL-1992) were cultured at 37°C and 5% CO2 in RPMI 1640 supplemented with 10% FCS, 10 mM HEPES, 100 U/mL penicillin, 100 µg/mL streptomycin, 1X non-essential amino acids and 1mM sodium pyruvate.

The full-length, codon-optimized AV23.1 and BV13.1 chain sequences of NY-ESO-1_157-165_-specific 1G4, 1G4-c50 and 1G4-c53c50 TCRs, separated by an IRES module, were synthetized by GeneScript and cloned into the MCS BamHi/XhoI of SFG retroviral vector. Additional amino acid substitutions within the CDR3β loops of 1G4, 1G4-c50 and 1G4-c53c50 TCRs were generated by introducing short 81bp mutagenic ssDNA fragments using the NEBuilder HiFi DNA Assembly protocol (Biolegend) according to the manufacturer’s instruction. Constructs were transformed into XL10-Gold ultracompetent bacteria (Agilent) and full-length TCRaβ sequences were confirmed by DNA sequencing.

Retroviral vectors were produced by transient transfection of 293T cells in 100 μL DMEM medium supplemented with 3% GeneJuice transfection reagent (Sigma-Aldrich) with the vector of interest (SFG.TCR AV23.1-IRES-TCR BV13.1) and the PegPam3 (gag-pol) and RDF (env) plasmids. Supernatant of retroviral-transfected 293T cells was used to transduce HLA-A*02:01^neg^ or HLA-A*02:01^pos^ J76 CD8αβ cells using RetroNectin (Takara) coated plates. TCR-positive HLA-A*02:01^neg^ or HLA-A*02:01^pos^ J76 CD8αβ cells were sorted to purity by flow cytometry (FACSAria II and III, BD Biosciences) using PE-labeled A2/NY-ESO-1_157-165-specific multimers (Peptide & Tetramer Core Facility, UNIL-CHUV Lausanne) and anti-Vβ13.1 APC antibodies (BioLegend).

### Functional assay - cross-reactivity assay

HLA-A*02:01^pos^ TAP-deficient T2 cells were loaded with three peptides (FLTLWLTQV, GLRMWIKQV and TQIQWATQV) from the self-proteome, with NY-ESO-1 (SLLMWITQA) or the CMV/pp65 (NLVPMVATV) peptide at 0.01, 0.1 or 1 µg/mL at 37°C for 1h. We used the heteroclitic SLLMWITQA peptide instead of SLLMWITQC to avoid disulfide bridge formation, improving loading onto HLA complexes and T cell responses ^35,85^. Peptide-loaded T2 targets were cocultured with TCR-transduced HLA-A*02:01^neg^ J76 CD8αβ cells at a 3:1 ratio (1.5x105 T2 and 0.5x105 J76) during 16h in U-bottom 96-well plates. Co-cultures using unloaded “empty” T2 targets or parental HLA-A*02:01^neg^ J76 cells without any TCR expression (no transduction) were used as additional negative controls, while TCR-transduced J76 cells stimulated with PMA (500 ng/mL) / Ionomycin (250 ng/mL) were used as additional positive control (Supplementary Figure 6).

### Functional assay - self-proteome assay

5x10^4^ freshly TCR-transduced HLA-A*02:01^pos^ J76 CD8αβ cells were cultured during 3 to 6 days in U-bottom 96-well plate under steady-state culture conditions for testing the presentation of endogenous epitopes derived from the self-proteome. Parental HLA-A*02:01^pos^ J76 cells without any TCR expression (no transduction) were used as negative controls while HLA-A*02:01^pos^ J76 cells expressing the high affinity 1G4-c53c50 TCR variant were used as an positive control (Figure 4).

### Functional assay - surface staining by flow cytometry

1x10^5^ to 2x10^5^ TCR-transduced HLA-A*02:01^pos^ J76 CD8αβ cells (from the self-proteome assay) or TCR-transduced HLA-A*02:01^neg^ J76 CD8αβ cells (from the cross-reactivity assay) were stained at room temperature with anti-CD69 PerCP-eF710 (Invitrogen), anti-PD-1 PE (BioLegend) and anti-Vβ13.1 APC (BioLegend) antibodies for 20 minutes. For the cross-reactivity assay, anti-CD20 FITC (BioLegend) was added to gate out the CD20+ peptide-loaded T2 target cells. DAPI was used as a dead cell marker. Samples were acquired on a Cytoflex (Beckman Coulter) flow cytometer and data were analyzed by FlowJo software (Tree star, v.10.8.1).

### Selection of peptides showing high similarity to NY-ESO-1

The human proteome data, excluding isoforms, was downloaded from UniProt (https://www.uniprot.org/proteomes/UP000005640), from which we generated and all possible 9-mers were extracted. MixMHCpred ^83^ was then used to predict the binding affinity of these peptides to HLA-A*02:01. Only cases having %rank values below 1.5 were selected for further analysis. Next, we computed the sequence similarity between a peptide and the NY-ESO-1 epitope (SLLMWITQC) using the BLOSUM62 scoring matrix ^86^ from the biopython package ^87^. Among the top scoring cases, we selected three peptides (FLTLWLTQV, GLRMWIKQV, and TQIQWATQV) having specific residues (W5, Q8) which are known to be critical for TCR binding. The FLTLWLTQV and TQIQWATQV peptides were also reported in an earlier study investigating the cross-reactivity of the affinity-enhanced TCR NY-ESOc259 ^37^.

### Structural analysis

Structural modeling of the TCR-pMHC complexes with modified CDR3β sequences was performed using the Modeller software ^88^ (version 10.2). The 2BNR crystal structure served as a template, with CDR3β sequences integrated into the TCR while maintaining the fixed coordinates of the pMHC and non-CDR3β TCR residues. Following this, the best 5 models were selected based on the DOPE score evaluated for the TCR-pMHC interface encompassing CDR loops, the peptide and MHC residues located within 6 Å from the peptide. Among the top five models, the one with the maximal number of hydrogen bonds and hydrophobic contacts was retained. The solvent accessibility of each CDR3β residue was determined as the relative solvent excluded surface area (SESA) computed with the MSMS package of the UCSF Chimera software ^89–91^. The normalized SESA, nSESA, was calculated by normalizing the surface area of the residue in the TCR of interest by its surface area in a reference state. The latter was defined as the Gly-X-Gly tripeptides in which X is the residue type of interest ^92^. nSESA thus ranges from 0% for totally buried residues to 100% for residues exposed to the solvent to the same degree as in Gly-X-Gly. The Beta factor of the core region of the CDR3β loop for the 1G4 template was obtained from the PDB entry 2BNR.

## Data availability

The raw sequencing data (fastq files) of the input and output phage-display libraries are stored at European Nucleotide Archive (ENA) under the project number PRJEB76298.

## Code availability

The pretrained MixTCRpred model A0201_NY-ESO-1-CDR3b is available at https://github.com/GfellerLab/MixTCRpred/tree/ny_eso_1_phage.

## Declaration of interests

David Gfeller is a consultant for CeCaVa and GNUbiotics. The remaining authors declare no competing interests.

## Acknowledgments

We thank Christophe Sauvage for his technical help. This project has received funding from the SNF Sinergia program (CRSII5_193749) to D.G., A.H., V.Z., M. AS P., M.B., and G.C; the European Union’s Horizon 2020 research and innovation program under the Marie Skłodowska-Curie grant agreement, No. 101027973 to G.C.; the KFS-4368-02-2018 grant to D.T., M.H., and N.R.; and the SNSF grant No. 205321_192019 to M. AS P., M.B., and V.Z. Figures 1-5 were created with Biorender.com.

## Contribution

D.G. and G.C. designed the study and wrote the paper. G.C. carried out the bioinformatics analyses. R.L., M.d.T. and SM. D. performed the phage display experiments. S.B., P.G., J.S. and A.H. conducted the multimer-sorting experiments. D.T., M.H. and N.R. conducted the functional assay. M.B., M. AS. P. and V.Z. conducted the structural analysis. All authors provided materials and feedback on the paper.

## Supplementary information

**Supplementary Figure 1.**
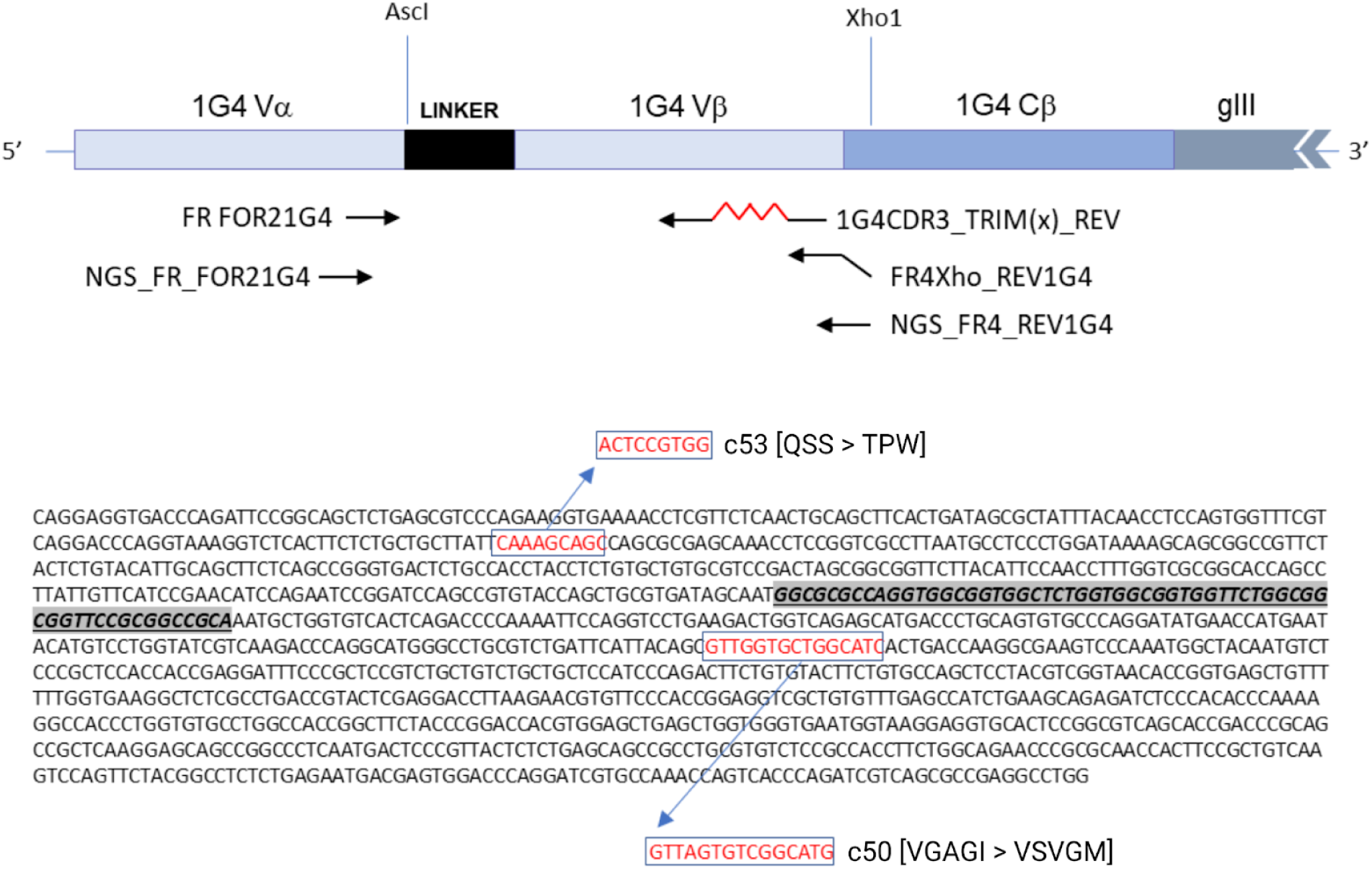
Schematic of 1G4 CDR3β PCR strategy (upper), and codon-optimized DNA sequence and variants (lower).

**Supplementary Figure 2.**
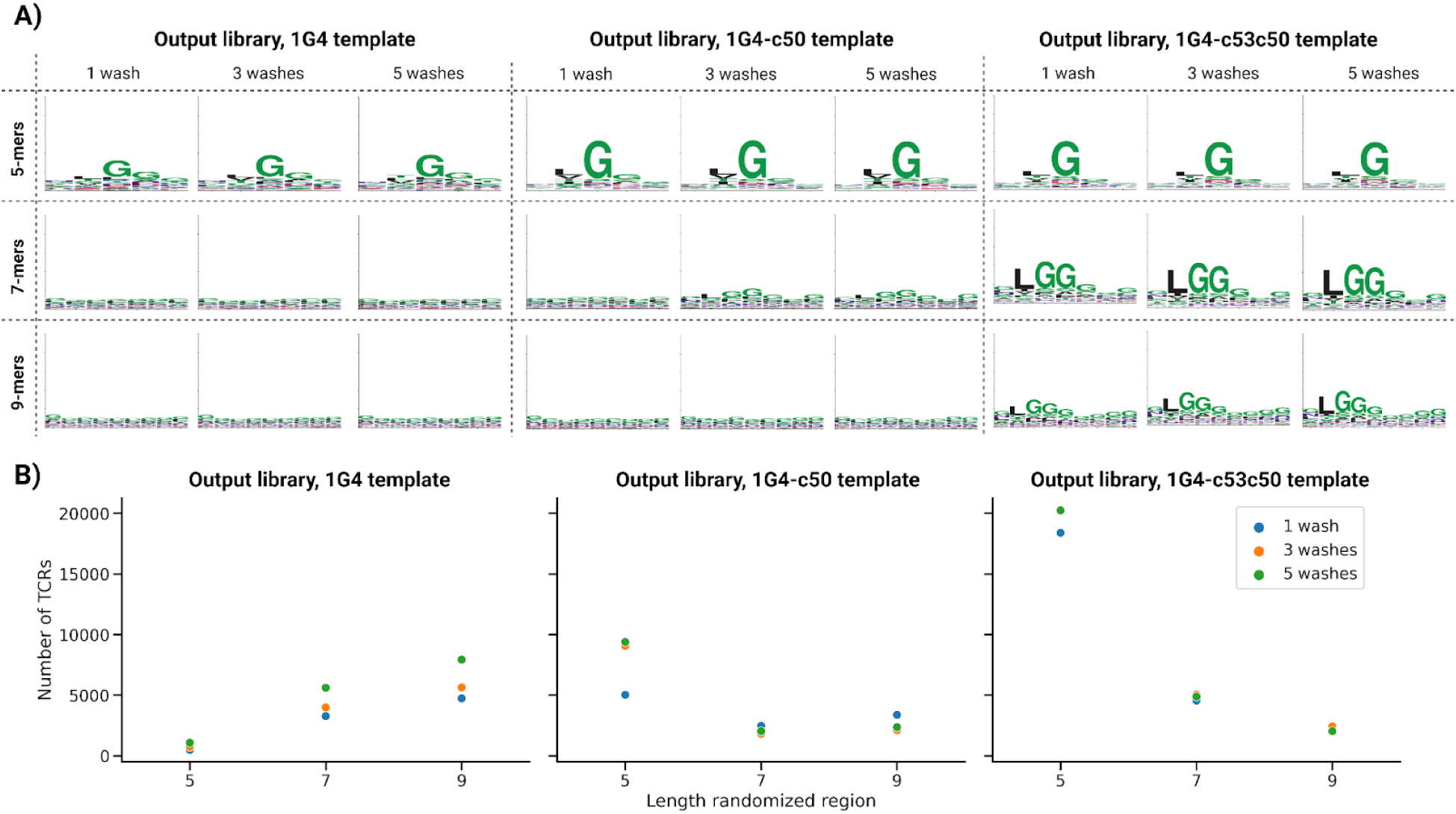
(A) Motifs of the raw TCR sequences resulting from the phage display screening with 1,3 and 5 washes. The data for each template TCR are shown separately. (B) Number of TCR sequences of different length resulting from the phage display screening with 1, 3 and 5 washes.

**Supplementary Figure 3.**
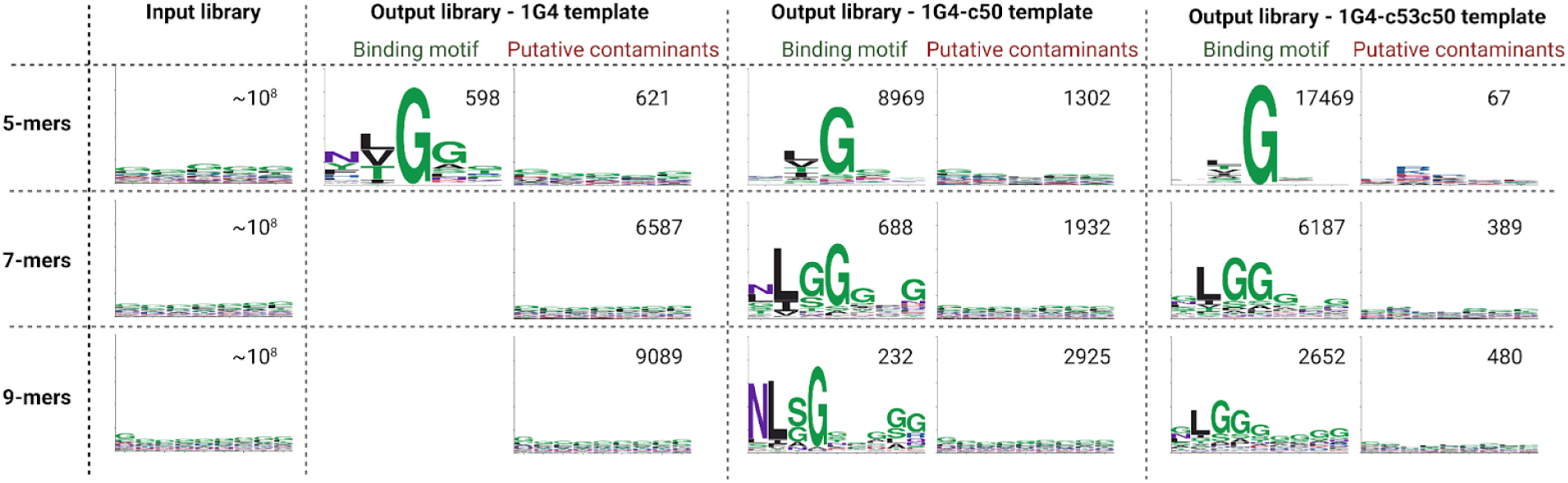
Motifs of the input and output libraries of the phage display screening after filtering out putative contaminants with the motif-deconvolution algorithm MoDec ^72^. For each motif, the number of TCR sequences is also reported.

**Supplementary Figure 4.**
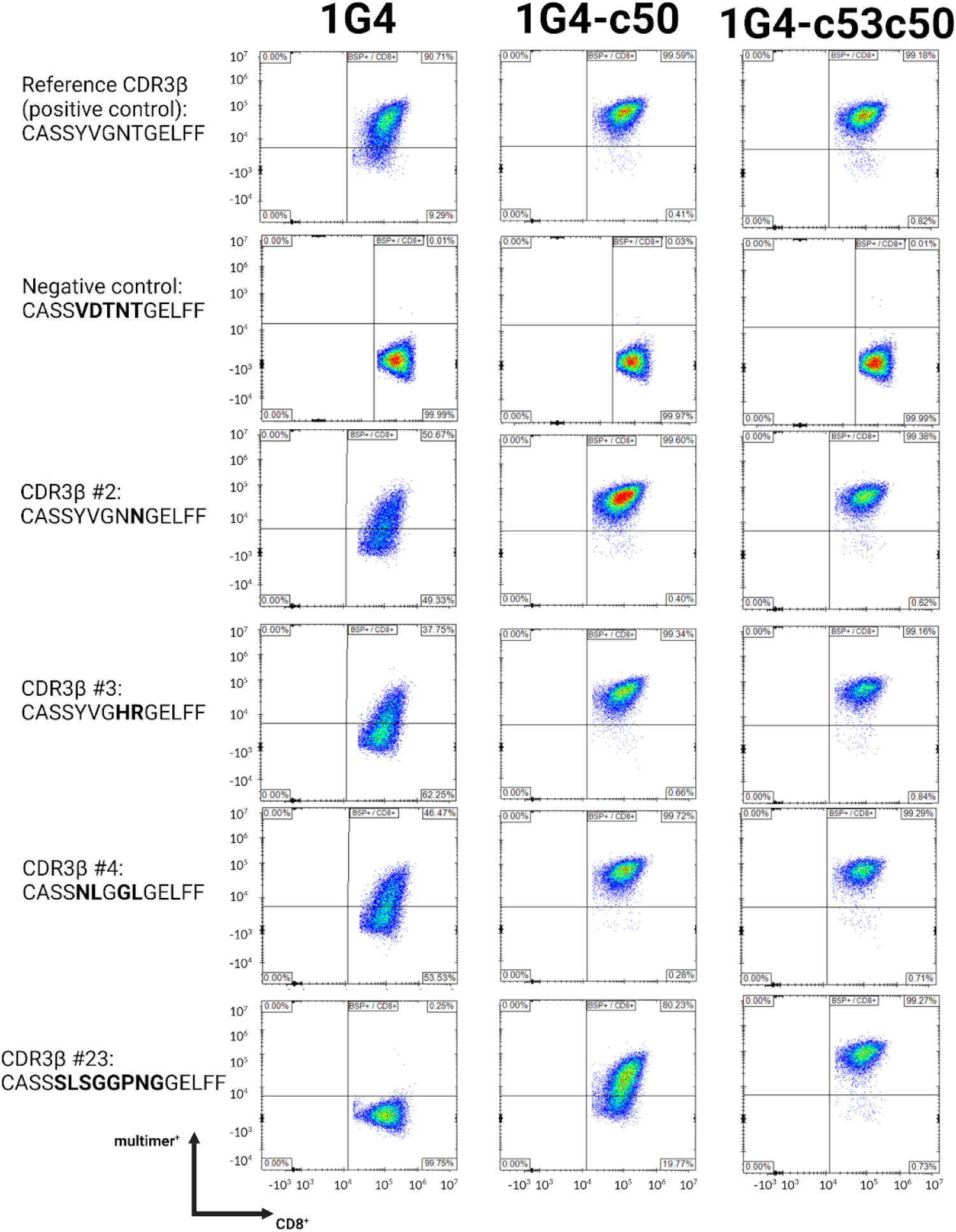
Representative FACS plots for the results of the multimer staining for five different CDR3β sequences.

**Supplementary Figure 5.**
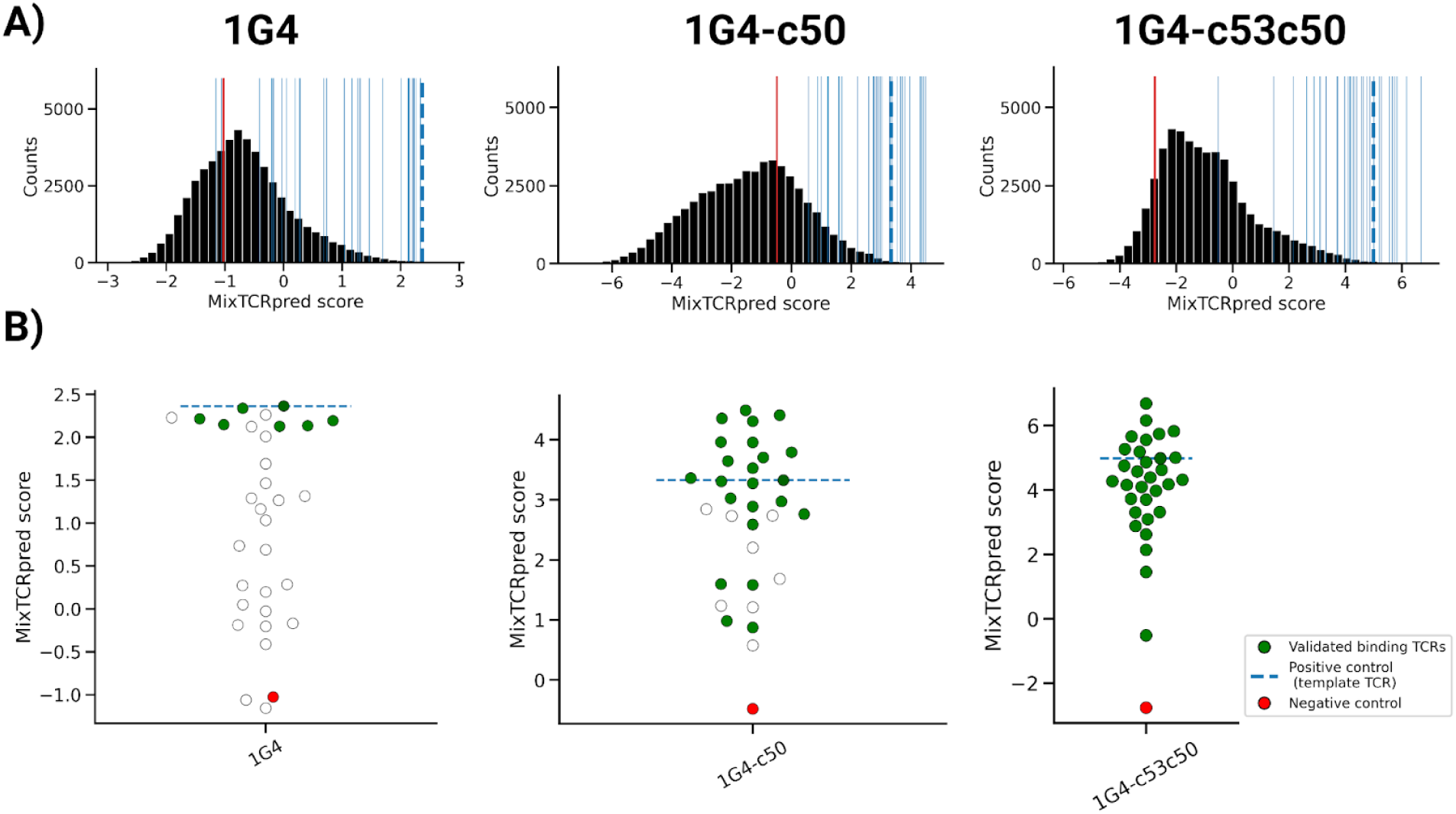
Results of the MixTCRpred models trained on data from each template separately. (A) Distribution of the MixTCRpred scores. The blue lines show the TCRs selected for experimental testing. The dashed blue line shows the template CDR3β (CASS**YVGNT**GELFF) while the red line shows the negative control (CASS**VDTNT**GELFF). (B) Scores of the CDR3β sequences that could be (green) or not be (white) experimentally validated on the 1G4, 1G4-c50, and 1G4-c53c50 templates. The negative control is shown in red.

**Supplementary Figure 6.**
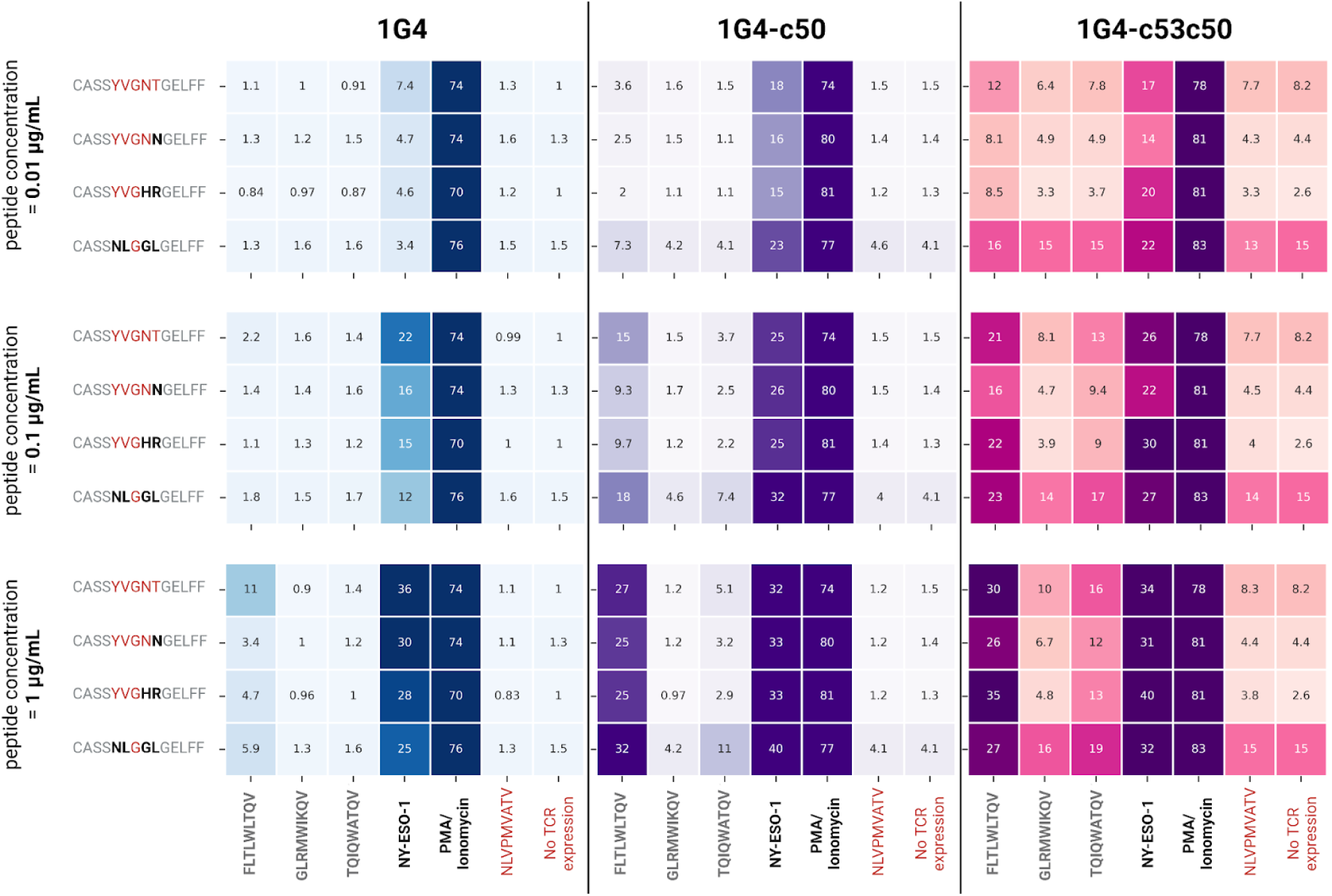
Heatmaps showing the fraction of CD69+PD-1+ Jurkat cells encoding four TCRs with different CDR3β sequences based on the three template TCRs. Jurkat cells were co-cultured overnight with peptide-pulsed T2 cells, at multiple peptide concentrations (0.01 μg/mL, 0.1 μg/mL, and 1 μg/mL).

**Supplementary Table 1.** Codon mixture of the diversified CDR3β library devised to approximate the composition of core regions of CDR3β loop in TCR repertoires.

| AA |  | Codon (antisense) | % in mixture |
| --- | --- | --- | --- |
| F | Phe | GAA | 2 |
| V | Val | AAC | 4 |
| D | Asp | ATC | 5 |
| H | His | ATG | 1 |
| L | Leu | CAG | 8 |
| M | Met | CAT | 1 |
| R | Arg | TCT | 7 |
| Q | Gln | CTG | 5 |
| S | Ser | AGA | 10 |
| T | Thr | GGT | 8 |
| N | Asn | GTT | 4 |
| A | Ala | TGC | 7 |
| P | Pro | TGG | 6 |
| E | Glu | TTC | 3 |
| I | Iso | GAT | 2 |
| W | Trp | CCA | 1 |
| G | Gly | ACC | 22 |
| K | Lys | TTT | 1 |
| Y | Tyr | GTA | 3 |

**Supplementary Table 2.**
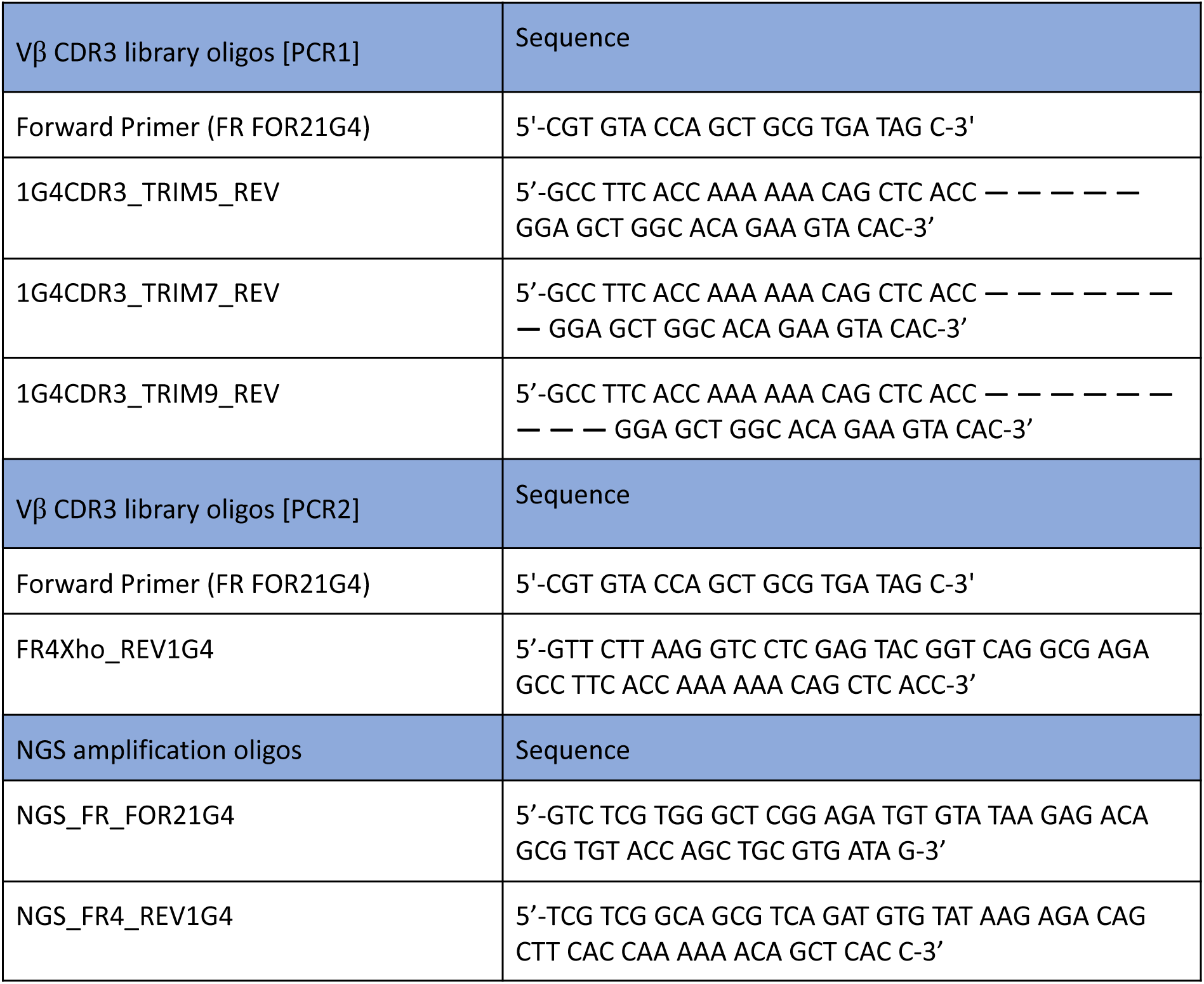
Primers used in the phage display screening.

**Supplementary Data 1.** The randomized CDR3β sequences of the input phage libraries.

**Supplementary Data 2.** The CDR3β sequences obtained with the phage display screening after panning with the NY-ESO-1 epitope, and the results of the motif deconvolution.

## Notes

### Summary of Updates

The title was changed; Abstract was modified to clarify the novelty of the approach

